# Effect of pharmacokinetically-relevant ivermectin concentrations on survivorship and fecundity of *Anopheles coluzzii* and *Aedes aegypti* in Burkina Faso: a laboratory experimental study

**DOI:** 10.1101/2025.09.04.674356

**Authors:** Emmanuel Sougué, Cheick Oumar W. Ouédraogo, Fabrice A. Somé, Greg Pugh, André B. Sagna, S. Rodrigue Dah, Saoudatou Bagayogo, Risnahar T. Ouédraogo, Mady Ndiaye, Sunil Parikh, El Hadji M. Niang, Brian D. Foy, Roch K. Dabiré

## Abstract

**Background:** The control of vector-borne diseases is increasingly challenged by widespread insecticide resistance. Therefore, innovative vector control strategies with alternative modes of action are urgently needed. Ivermectin (IVM), an endectocide, has demonstrated toxicity to mosquito species such as *Anopheles* and *Aedes* when they feed on treated humans or livestock. In this study, we conducted a laboratory experiment to assess the effect of IVM, at concentrations equivalent to human plasma levels following mass drug administration (MDA), on the survival and fecundity of *Anopheles coluzzii* and *Aedes aegypti* in Burkina Faso.

**Methods:** Two laboratory experiments were conducted using 3–5-day old wild-derived female *An. coluzzii* and *Aedes aegypti*. Each experiment included four replicates per IVM concentration and was performed on separate dates. Mosquitoes were fed via membrane feeding on rabbit blood treated with five concentrations of IVM (C=112 ng/ml, C2=29 ng/ml, C3=15 ng/ml, C4=6.5 ng/ml, C5=2.5ng/ml), corresponding to the mean human plasma levels at 2, 4, 7, 14, and 28 days post-MDA with IVM at a dose of 300 μg/kg. A negative control (C6=0.0 ng/ml) was also included. Mosquito mortalities were recorded daily for 7 days. Fecundity was measured by counting both laid eggs and developed eggs (via ovary dissection).

**Results:** IVM significantly reduced the survival of *An. coluzzii* compared to the control group (*p<0*.*001*), with the risk of death increasing from 4.2-fold at the lowest concentration (2.5 ng/ml) to 64.2-fold at the highest (112 ng/ml). In contrast, IVM had no significant effect on *Aedes aegypti* (*p*>*0*.*05*). Additionally, in *An. coluzzii*, IVM significantly reduced both egg laying and egg development (p<0.0001 and p<0.001, respectively), whereas no significant impact on fecundity was observed in *Ae. aegypti* (all *p>0*.*80)*.

**Conclusion:** Ivermectin concentrations typically achieved in human plasma during mass drug administration campaigns were sufficient to significantly reduce both survival and fecundity of wild type *An. coluzzii*, but had no measurable effect on recently colonized *Ae. aegypti*. These findings highlight the species-specific efficacy of ivermectin and support its potential role in integrated vector control strategies targeting malaria vectors in Africa.

## Introduction

Vector-borne diseases such as malaria and dengue remain major public health challenges, affecting millions of people worldwide each year. In 2023, the World Health Organization (WHO) estimated approximatively 263 million malaria cases and approximately 597,000 related deaths, with about 95% of fatalities occurring in African countries, where many at-risk populations still lack access to preventive measures [1]. In Burkina Faso, approximately 8,139,000 malaria cases and 16,146 deaths were recorded in the same year [1]. Dengue fever also surged in 2023, with WHO reporting a record five million cases and over 5,000 deaths worldwide. In Africa, Burkina Faso was the most affected country during this outbreak with 146,878 suspected cases, 68,346 confirmed by Rapid Diagnostic Test (RDT) and 688 associated deaths [1].

In regions where malaria and dengue are co-endemic, vector control strategies primarily rely essentially on indoor residual spraying (IRS),long-lasting insecticide-treated bed nets (LLINs) and larval source management (LSM) [2]. These interventions utilize insecticides, namely pyrethroids, that target the voltage-gated sodium channels in mosquito neurons. However, the effectiveness of these tools is increasingly compromised by widespread insecticide resistance [3] and behavioral changes, including shifts in biting times (earlier or later), biting and resting outside of human dwellings (exophagy and exophily), and feeding preference (zoophagy) [4-6]. This growing challenge of multi-factorial resistance and the continued propagation of vector borne diseases around the world [7] highlight the urgent need for novel control tools that can target mosquitoes regardless of their biting or resting behavior, including those that feed outdoors or during daytime hours [8].

Ivermectin (IVM), a semi-synthetic derivative of avermectin, was approved for human use in 1987 [9]. It is primarily used to treat parasitic diseases such as onchocerciasis, lymphatic filariasis headlice, strongyloidiasis and scabies [10, 11]. IVM is well tolerated in both humans and animals and is widely used in mass drug administration (MDA) campaigns for controlling parasitic infections such as onchocerciasis and lymphatic filariasis. It is also routinely administered to livestock to manage helminth infections [12]. Beyond its anti-parasitic properties, IVM has demonstrated mosquitocidal effect in several studies [12-16], leading to its consideration as a novel strategy for vector control [17-21]. Indeed, unlike conventional public health insecticides, IVM targets the glutamate-gated chloride channels in mosquito nerve and muscle cells, leading to an uncontrolled influx of chloride ions, resulting in paralysis and death [22, 23]. The recommended human dose during MDA is 150 mg/kg body weight, typically resulting in peak plasma concentrations of 40-45 ng/ml [24]. These levels have been shown to reduce survival rates of *Anopheles* mosquitoes for 5-7 days post-treatment [25-27]. Due to it safety profile, higher and more sustained plasma concentration can be achieved by frequently administering IVM at a 300μg/kg dose [28].

While studies have evaluated the impact of IVM on the survival of *Anopheles* and *Aedes* mosquitoes, most used laboratory-colonized strains. Its effects on wild-type *Anopheles* and *Aedes* remain understudied; hence these vectors are often sympatric and feed on the same human hosts [29]. In this study, a laboratory experiment was performed to investigate the effects of IVM on survival and fecundity of wild-type *Anopheles coluzzii* and recently-colonized *Aedes aegypti*, which are sympatric and the primary vectors of malaria and dengue in most of Burkina Faso. Mosquitoes were membrane fed with blood treat with IVM at that mimic human plasma levels at different timepoints following high dose MDAs.

## Materials and methods

### Mosquito rearing and maintenance

Gravid *An. coluzzii* females were collected from human dwellings in Bama (11°23’14’’N,4°24’42’’W) in January and May 2024. They were placed individually in cups containing water and covered with mesh netting to allow for oviposition. Species were confirmed by routine PCR [30] after oviposition. All *An. coluzzii* larvae were pooled and reared in tap water and fed with Tetramin® Baby Fish food (Tetrawerke, Melle, Germany). *Aedes aegypti* used for the experiments was a wild-derived, originating from larvae collected in breeding sites around the city of Bobo-Dioulasso (11°10’37’’N, 4°14’52’’W) in October 2022.

All mosquitoes were maintained in 30×30×30 cm cages covered by mosquito netting, provided with 10% glucose solution, and reared under standard insectary conditions (temperature: 27±2°C; relative humidity: 70±10%; photoperiod: 12h light followed by 12h dark).

### Study drug preparation

Powdered IVM was obtained from Sigma-Aldrich (St. Louis, MO, USA). A stock solution was prepared at a concentration of 10 mg/ml in dimethyl sulfoxide (DMSO) and refrigerated overnight at 4°C. This stock solution was then diluted in DMSO to create three working solutions of 1 mg/ml, 0.1 mg/ml and 0.01 mg/ml (**Supplemental file Fig 1**). From the 0.01mg/ml solution, five concentrations (C1= 112 ng/mml, C2= 29 ng/ml, C3= 15 ng/ml, C4 =6.5 ng/ml, and C5= 2.5 ng/ml) corresponding to mean IVM plasma concentrations measured at days 2, 4, 7, 14, and 28 days, respectively, following MDA with IVM. The values were based on pharmacokinetic/pharmacodynamic (PK/PD) data from a sub-study of the RIMDAMAL II trial [31]. Each concentration was mixed with rabbit blood for use in mosquito bioassays.

### Mosquito blood-feeding assays

Blood-feeding assays were conducted using freshly collected rabbit blood on each experimental day. For both mosquito species, two independent experiments were conducted on different dates, with four replicates per IVM concentration. In each experiment and for each strain, four test cups containing approximately 50 females aged 3–5 days-old were used. Females were starved of sugar for 8 hours prior to feeding assays. Blood meals were offered using Hemoteck membrane feeders (Hemotek Ltd., UK), covered with parafilm, heated to 37°C, and placed on top of each cup. Mosquitoes were exposed to IVM-treated blood for 30 minutes (**Fig 1**). After feeding, unfed and partially blood fed mosquitoes were discarded. Fully-engorged mosquitoes were retained, allowed to recover, and maintained in the insectary at 27-29°C, ∼70% relative humidity (RH) with access to water and 10% sucrose for 7 days.

**Fig 1:**
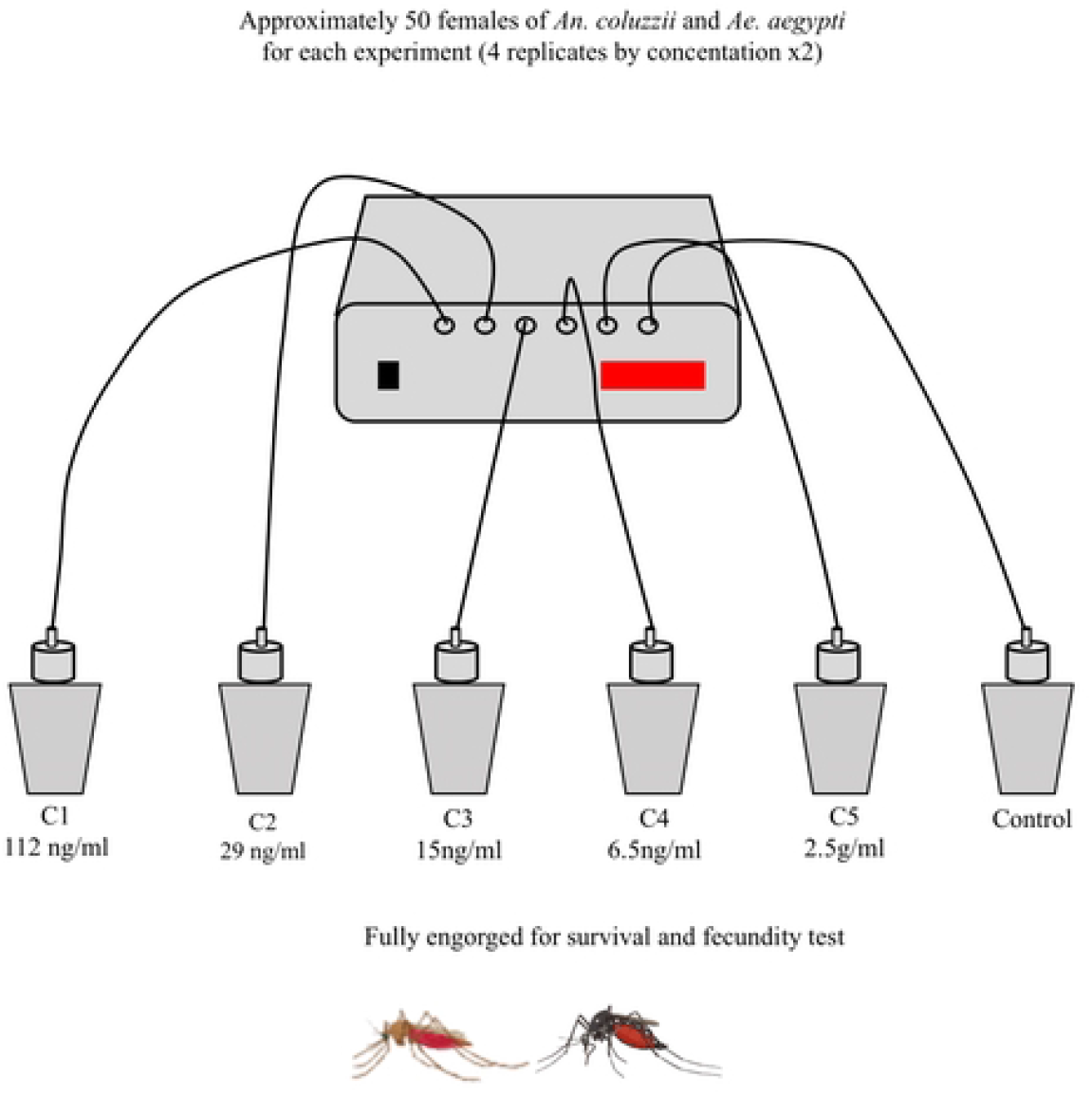
Experimental workflow.

### Mosquito survival and fecundity assays

Mosquito survival was monitored daily, and dead mosquitoes were removed. Mosquitoes were considered dead if unable to stand on their legs.

For fecundity assessments, a similar feeding setup was used. Approximatively 50 unfed females were fed with blood containing different ivermectin concentrations (four replicates per concentration). Fecundity assays started 3 days post blood feeding, due to high mortality during the first two days post feeding. Only surviving mosquitoes were kept in labelled cups, where petri dishes containing moist cotton and filter paper were provided as oviposition support for gravid females. Mosquitoes were maintained for five days, with access to 10% glucose solution. Eggs laid on filter paper were counted under binocular microscope at 2X magnification. Afterward, all mosquitoes were dissected to examine their ovaries and count any undeveloped or retained eggs that were not laid.

### Statistical analysis

Data were entered and cleaned using Microsoft Excel 2021, and statistical analyses were performed using R software version 4.4.1. Mosquito survival data for each species, were visualized using Kaplan-Meier curves, and differences among IVM concentrations were assessed using the Log-rank Mantel-Cox test and the Kruskal-Wallis non-parametric ANOVA. Lethal concentrations (LC values) were estimated using non-linear regression analyses with the *drc* package for R software [32]. Generalized linear regression models were employed to assess the impact of ivermectin on egg production for surviving female mosquitoes. The mean number of developed and/or laid eggs per female according ivermectin concentration was assess using a correlation analysis. All *p* values <0.05 were considered statistically significant and retained in the minimal adequate model [33].

## Results

### Effect of ivermectin on mosquito survival

A total of 2,400 *An. coluzzii* and 1,200 *Ae. aegypti* females were used to assess the effect of IVM on mosquito survival. Following blood feeding, 1,290 *An. coluzzii* and 548 *Ae. aegypti* were fully engorged and used for survival assays.

IVM concentration had no significant effect on blood-feeding rates in either species. For *An. coluzzii*, blood-feeding rates were 47%, 51.5%, 53.5%, 52.2%, 52.7%, and 67% for concentration C1 (112 ng/ml), C2 (29 ng/ml), C3 (15 ng/ml), C4 (6.5 ng/ml), C5 (2.5 ng/ml), and control, respectively. For *Ae. Aegypti*, blood-feeding rates were 39.25%, 46.5%, and 51.25 for C1 (112ng/ml), C2 (29ng/ml) and control.

Survival probabilities of both species over 7 days post-ingestion of blood containing different IVM concentrations are shown in **Fig 2**. IVM had no significant impact on the survival of *Ae. aegypti* (*p>0*.*05*). On day 7 post-ingestion, survival remained at 100% in the control group and approximately 95% for the two concentrations tested (C1 and C2; **Fig 2)**. As a result, LCs and HRs could not be estimated for this species, and the experiments with *Ae. Aegypti* was discontinued.

**Fig 2:**
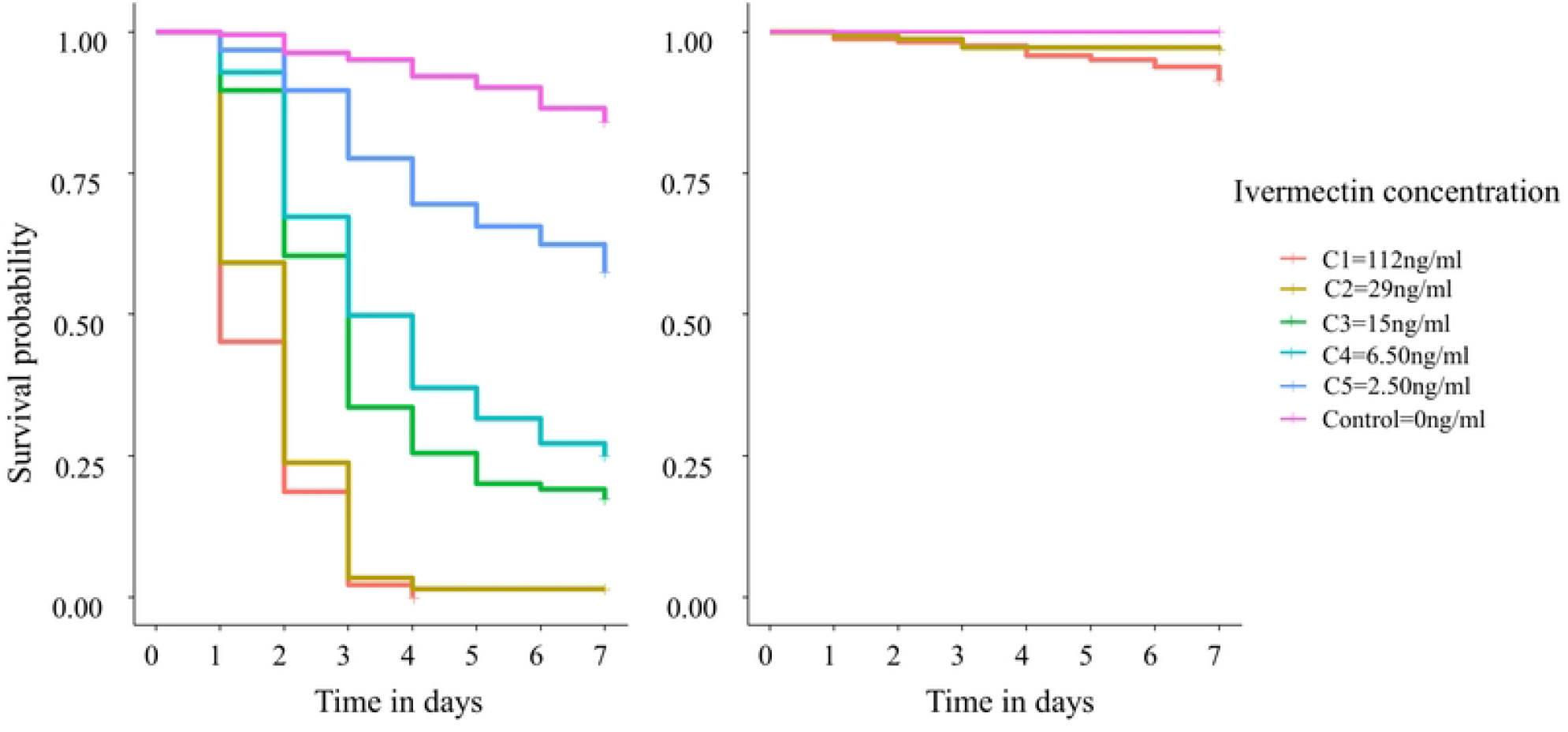
Kaplan-Meier 7 days survival probability of *An. coluzzii* (left) and *Ae. aegypti* (right) according to ivermectin concentration.

In contrast, *An. coluzzii* showed high survival in the control group, while survival was significantly reduced in all IVM-treated groups (LRT χ^2^ = 960.9, df = 5, *p<* 0.001, **Fig 2**). One day after blood feeding, survival was reduced by 50% with C1 (112ng/ml), 35% with C2 (29 ng/ml), 13% with C3 (15 ng/ml), 7% with C4 (6.50 ng/ml) and 3% with C5 (2.50 ng/ml). By day 3 post-feeding, survival dropped further to reach approximatively 5% for both C1 (112ng/ml) and C2 (29ng/ml), 35% for C3 (15 ng/ml), 50% for C4 (6.50 ng/ml), and 76.56% for C5 (2.50 ng/ml). Survival fell to 0% for C1 (112 ng/ml) by day 4, while remaining stable at approximatively 5% until day 7 post-feeding for C2 (29 ng/ml). *An. coluzzii* survival continued to drop until day 7 post-ingestion for the 3 remaining concentrations to reach 23% for C3 (15 ng/ml), 28% for C4 (6.50 ng/ml), and 65% C5 (2.50 ng/ml).

All IVM concentrations had a statistically significant negative effect on *An. coluzzii* survival when compared individually to the control group. Mosquitoes that fed on IVM-treated blood were more likely to die compared to the control group (**Fig 3**). The 7-days HRs were ≥50 for C1 (112 ng/ml) and C2 (29 ng/ml), between 10 and 20 for C3 (15 ng/ml) and C4 (6.50 ng/ml), and below 10 for C5 (2.50 ng/ml).

**Fig 3:**
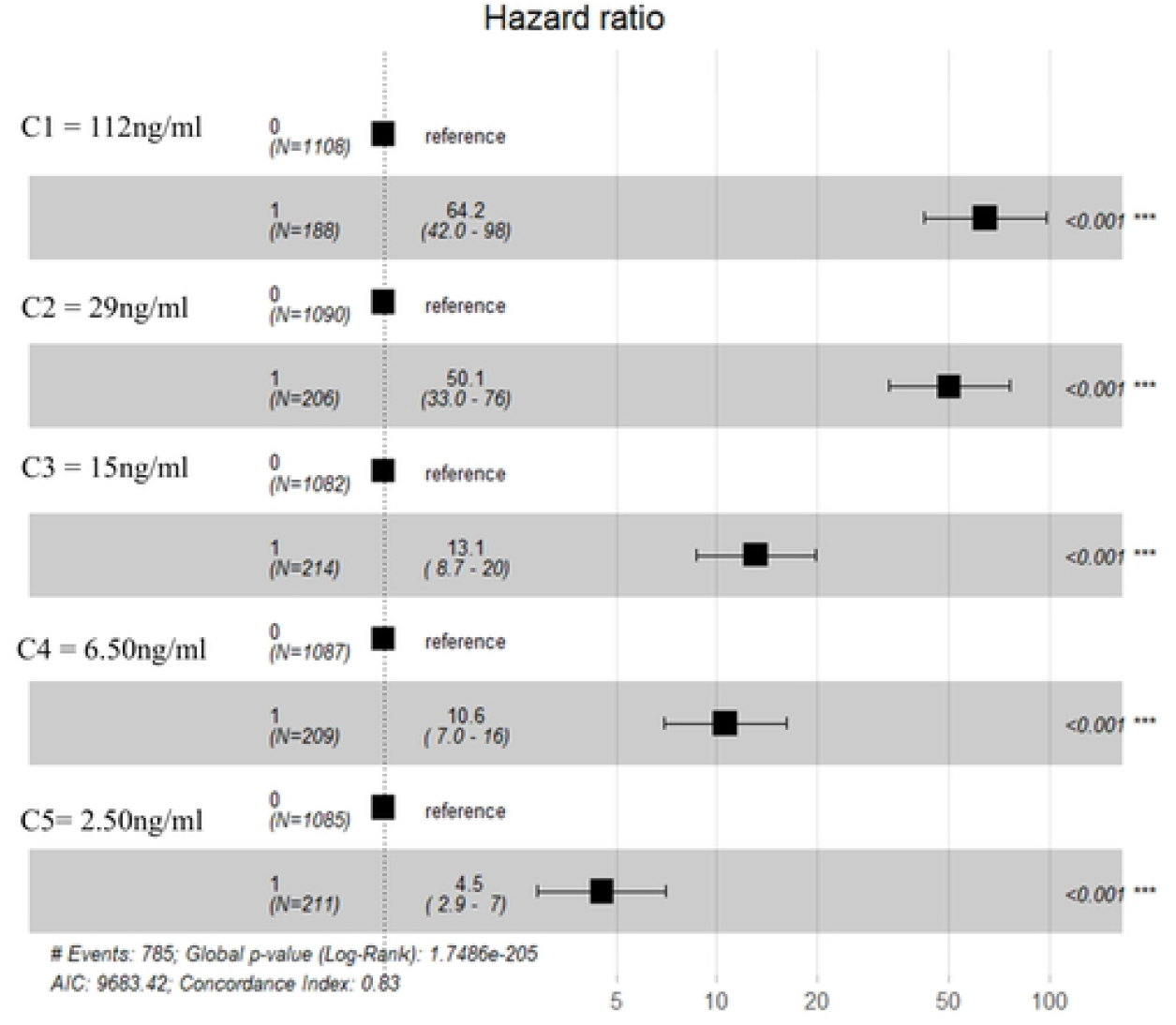
7-day Hazard Ratio estimations for *An. Coluzzii*.

Dose-response curves at days 1, 3, and 7 are presented in **supplemental file Fig 2**, illustrating cumulative mortality across different IVM concentrations during the 7-day observation period. Estimated lethal concentrations (LCs) resulting in 20%, 50%, and 90% mortality are summarized in Table 1. The LC50 (lethal concentration for 50% mortality) varied depending on the time point used for calculation. At day 1 post-ingestion, the LC50 was 75.2 ng/ml. When calculated over three days, it dropped to 9.4 ng/ml, and further decreased to 7.1 ng/ml by day 7 (**Table 1**).

**Table 1:** Lethal concentrations (LC) of ivermectin to *An. coluzzii* calculated at days 1, 3 and 7 post-blood feeding with their 95% CI.

| LC (%) | Day 1 | Day 3 | Day 7 |
| --- | --- | --- | --- |
| 20 | 20.84 [17.623 – 24.44] | 3.85 [3.11 – 4.59] | 1.92 [1.38 – 2.45] |
| 50 | 75.18 [59.80 – 90.57] | 9.44 [8.31 – 10.56] | 7.07 [5.40 – 8.74] |
| 90 | 574.16 [312.61 – 836.61] | 39.10 [31.43 – 46.99] | 55.87 [24.18 – 87.56] |

LC= Lethal concentration

### Effect of ivermectin on mosquitoes’ fecundity

Fecundity could not be assessed for *An. coluzzii* at IVM concentrations C1, C2, and C3 due to high post-feeding mortality. Exposure of *An. coluzzii* to IVM (C4 and C5) significantly reduced both the number of eggs laid (F = 32.14, df = 2, p < 0.001) and the number of developed eggs in the abdomen (F = 7.41, df = 2, p < 0.001). The mean number of laid eggs was 11.5 for C4, 57.2 for C5, and 105.1 for the control group. Similarly, the mean number of developed eggs in the abdomen was 50.6, 54.7, and 62.6 for C4, C5, and the control group, respectively (**Table 2**).

**Table 2:**
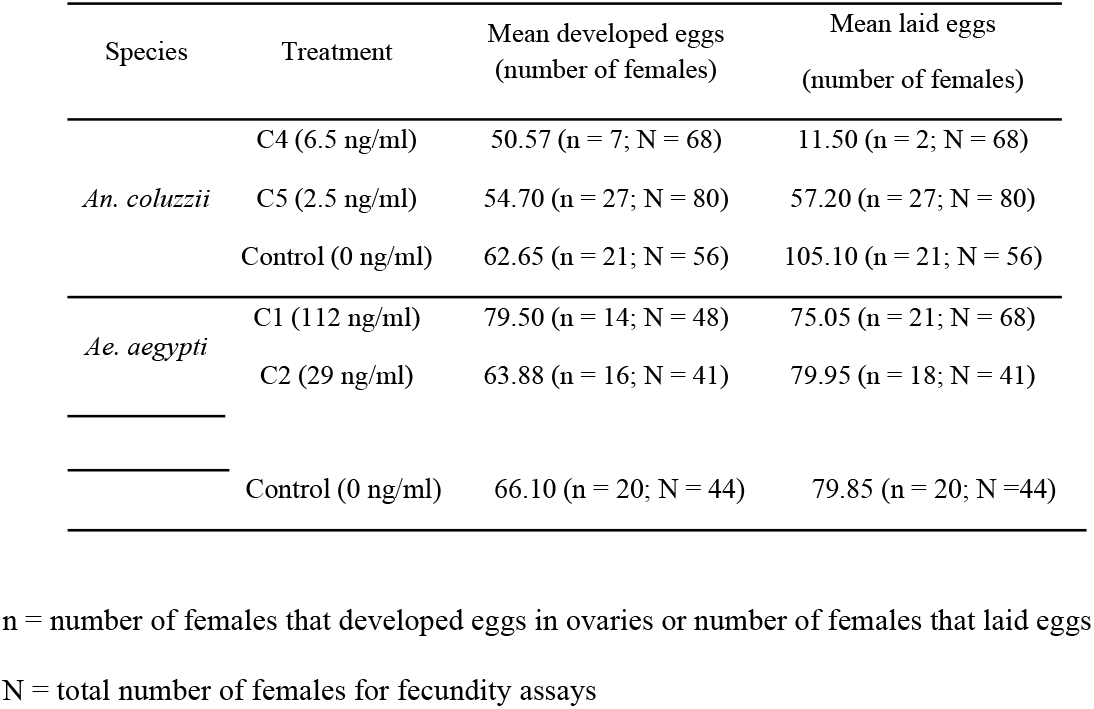
Fecundity of surviving *An. coluzzii* and *Ae. aegypti* females.

For *Ae. aegypti*, fecundity assays were only conducted at C1 and C2 as these high concentrations had shown no significant impact on survival. Unlike *An. coluzzii*, fecundity of surviving female *Ae. aegypti* was not significantly affected by IVM exposure. There were no significant differences in the number of eggs laid (F=32.14, df=2, p=0.88) or the number of developed eggs in the abdomen (F=0.16, df=2, p=0.85). The mean number of laid eggs was 75.1; 79.9, and 79.8 for C1, C2, and the control group, respectively. The mean number of developed eggs in the abdomen were 79.5 for C1, 63.9 for C2, and 66.1 for the control group (**Table 2**).

## Discussion

This study evaluated the impact of ivermectin (IVM), at concentrations equivalent to human plasma levels following mass drug administrations (MDA), on the survival and fecundity of *Anopheles coluzzii* and *Aedes aegypti* in Burkina Faso. Unlike many previous studies that used laboratory-susceptible mosquito strains [27, 34-40],. Unlike many previous studies that used laboratory-susceptible mosquito strains, the present study used derived F1 wild-type and recently colonized mosquito for experiments. we employed wild-type *An. coluzzii* and recently-colonized *Ae. aegypti* mosquitoes known to be resistant to pyrethroids [41-43]. IVM was delivered via membrane feeding using treated rabbit blood, mimicking plasma concentrations observed in human blood at 2, 4, 7, 14, and 28 days post-MDA with IVM at a dose of 300μg/kg (43). Our findings demonstrate a clear species-specific difference in IVM susceptibility. All tested IVM concentrations significantly reduced the survival and fecundity of female *An. coluzzii*, while no significant effect in both parameters was found for *Ae. Aegypti*.

The survival of *An. coluzzii* mosquitoes decreased to 50% at day 1 post IVM ingestion with an LC50 of 75.18 ng/ml. These results are comparable to those reported in a previous study [40]. Another study also revealed that when *An. arabiensis* mosquitoes blood fed on IVM-treated humans 1 to 4 days after treatment, their survival was significantly reduced [44]. Our study showed that the impact of IVM on mosquitoes is dose and species dependent. These results corroborate findings from previous studies demonstrating differential susceptibility of different mosquito species to ivermectin [27, 39]. The survival of wild type *An. coluzzii* mosquitoes in our study decreased significantly when they fed on blood containing IVM with all tested concentrations, and seven-day LC_50_ dose was 7.07ng/ml, in concordance with previous studies where seven-day LC_50_ for *An. gambiae* ranged between 3.35 to 55.60ng/ml [36, 37, 39, 45]. When mosquitoes blood fed on C1(112ng/ml) in our study, the majority died within 3 days, and practically all mosquitoes died on day 7 post exposure. The same trend was reported by previous studies with more than 90% of blood fed females dying by day 6 after ingestion of blood taken from IVM treated patients soon after treatment [34]. When *An. coluzzii* mosquitoes fed on IVM-treated blood groups C2=29ng/ml, C3=15ng/ml, and C4=6.50ng/ml corresponding to the concentrations found on days 4, 7 and 14 post-dosing with 300μg/kg, respectively, more than 50% died within 7 days. These results are similar to previous findings which Foley, Bryan (35) reported that *An. farauti* survival decreased significantly after day 14 when they blood fed on treated humans. Also, a recent field study conducted in Burkina Faso showed that survival rate of mosquitoes collected in villages one week after MDA with IVM decreased comparatively to those collected placebo-treated villages [31]. However, results derived from wild-type mosquitoes who blood fed directly on IVM treated human and collected directly from the field may be subject to bias, as these mosquitoes are often relatively old and may die from natural causes rather than as a result of IVM exposure. Additionally, field-collected mosquitoes may contain in their bodies, microorganisms such as parasites, viruses or bacteria like *Wolbachia*, which could have an impact on mosquitoes survival [46-48]. In contrast, the F1-generation mosquitoes we used were of known age, thus their survival rates can be consequently attributable to IVM. Furthermore, we observed that concentration 2.5ng/ml, corresponding to the IVM plasma concentration on the 28^th^ day after treatment with high dose IVM MDA, remained marginally but significantly lethal for *An. coluzzii*. Indeed, at this concentration about 40% of blood fed females died. However, a lethal effect did not detect 3 weeks after MDA they performed in their cluster randomized field trial, which may have been due to the diversity of the pharmacokinetic profiles among the thousands of participants treated, and the differences in how the mosquitoes were captured and tested in that trial relative to this lab-based study [49]. Although the lethal effect observed in the field seems to dissipate by the 28th day post dosing, IVM remains a complementary means for vector control as long as the life span of mosquitoes is reduced to prevent them from taking another blood meal and potentially transmitting malaria parasites [13].

Contrary to *An. coluzzii, Ae. aegypti* displayed a stronger tolerance to all IVM tested concentrations. Survival of *Ae. aegypti* did not decreased significantly after feeding on blood containing IVM at any of the tested concentrations, which reflects data from previous studies suggesting higher IVM concentrations are required to induce lethal and sublethal effect on *Ae. aegypti* [21, 50]. This tolerance may be due to the fact that IVM binds less effectively and/or has less stimulating effect on glutamate-activated channels in *Ae. aegypti* compared to *An. coluzzii* [27]. A previous study suggested that the relative tolerance of some mosquitoes species to IVM could be attributed to at least three reasons: 1) there may be poor absorption of IVM throughout the midgut, 2) there may be physiological differences in glutamate-gated chloride channel structures among mosquitoes species, and 3) some species may be more efficient in their ability to detoxify IVM compared to others [27]. Understanding this tolerance may be particularly critical for further development of endectocides like IVM to help control species like *Ae. aegypti* and limit their spread of arboviral diseases.

Our investigation of the fecundity of surviving female *An. coluzzii* after feeding on IVM treated blood revealed a significant reduction of the number of laid and developed eggs compared to the control group. In contrast, there was no significant impact on the fecundity of surviving female *Ae. aegypti*. This sub-lethal effect on *An. coluzzii* is in line with previous findings [51-54] suggesting a potential additional impact of ivermectin on the reduction of *Anopheles* density in the field by the reducing the fecundity of surviving females. In this study, our highest IVM concentration (C1=112ng/ml) did not impacted the fecundity of *Ae aegypti*. This result is consistent with a previous study that showed only concentrations ≥250ng/ml are able to induce an impact on *Aedes* fecundity [21].

The minor limitation of the present study is that all experiments were performed with diluted chemical IVM mixed with blood instead of blood directly sample on treated individuals during MDAs. Nonetheless, there are key contributions of this study to this important research topic. Firstly, its takes an integrated approach by simultaneously testing the major vectors of malaria parasites, *An. coluzzii*, and dengue viruses, *A. aegypti*. Secondly, this study used F1 mosquitoes derived from wild-type strains with a proven resistance status to pyrethroids. Finally, the concentrations tested were based on IVM plasma levels measured in humans following high-dose (300 μg/kg) MDA of IVM from a recent cluster-randomized trial [49].

## Conclusion

This study demonstrates that mean plasma concentrations of IVM achieved following high dose MDAs can induce high mortality and reduced fecundity in wild type *An. coluzzii* mosquitoes. While this effect decreased over concentrations that would be expected in humans over time post dosing, the effect was maintained at a mean plasma concentration that may be expected at days 28 post-MDA in some treated individuals. In addition to mortality, this study also showed that the fecundity of surviving *An. coluzzii* females was considerably reduced even at the lowest concentration tested. However, the human ivermectin plasma concentrations tested did not impact either mortality or fecundity in *Ae. aegypti* mosquitoes. Therefore, standard IVM drug formulations used in MDAs may be efficacious in controlling malaria spread by *An. coluzzii* in Burkina Faso, but these MDA could not be expected to simultaneously control dengue transmission by *Ae. aegypti* in the same treated regions. Further studies combining human MDAs using different combinations of endectocides, as well as livestock treatment, perhaps both with long lasting IVM formulation chemistries, could be more beneficial to achieve these goals. Regardless, past and current results testing IVM against certain mosquito vectors offer promise for vector-borne disease control, and continued research and careful implementation of studies and control interventions will be crucial to maximize its benefits while minimizing potential drawbacks.

## Availability of data and materials

Data are fully available from the corresponding author upon request.

## Authors’ contributions

ES, COWO, FAS, BDF and RKD conceived and designed the study. ES, COWO, SRD, RTO and SB collected the data. ES and COWO analyzed the data. ES drafted the original manuscript. COWO, GP, FAS, ABS, MN, SP, EHAMN, BDF, RKD reviewed the manuscript. All authors read and approved the final manuscript.

## Ethics approval

The study protocol has been approved by the institutional ethical committee with the CERS reference number 2019-01-009

## Consent for publication

Not applicable

---

**Supplemental Fig 1:**
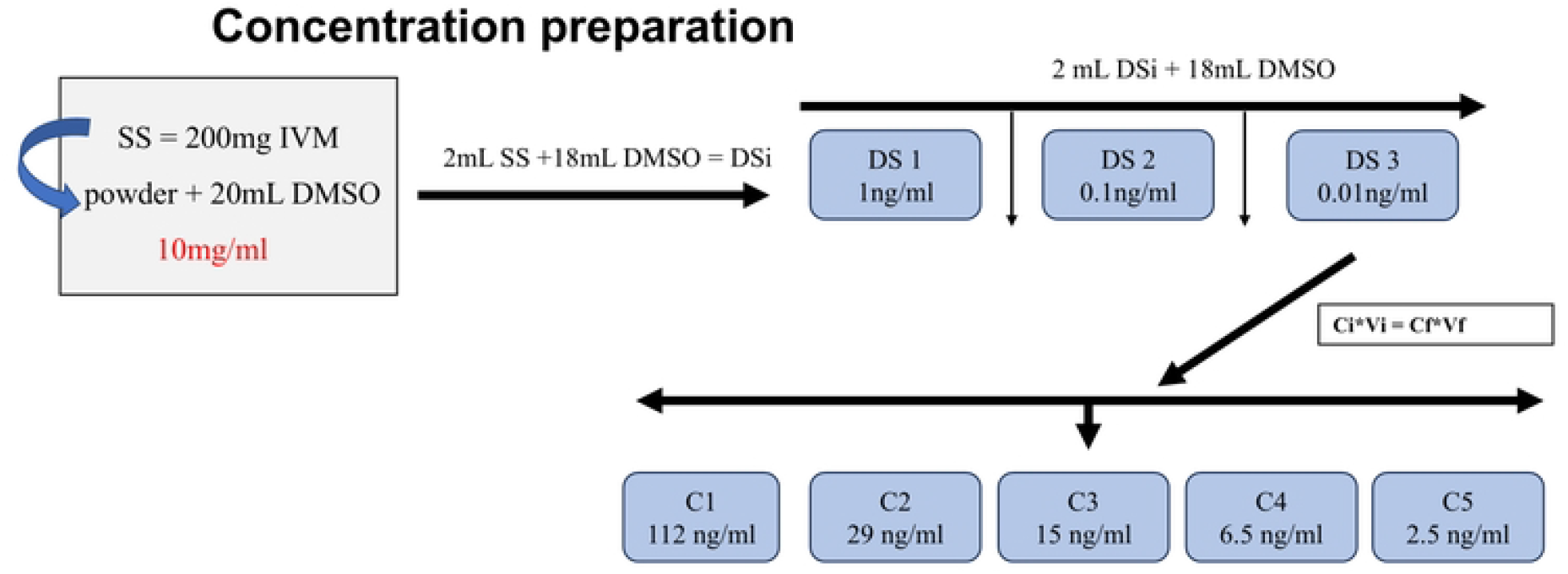
Ivemectin dilution and concentration preparation.

**Supplemental Fig 2:**
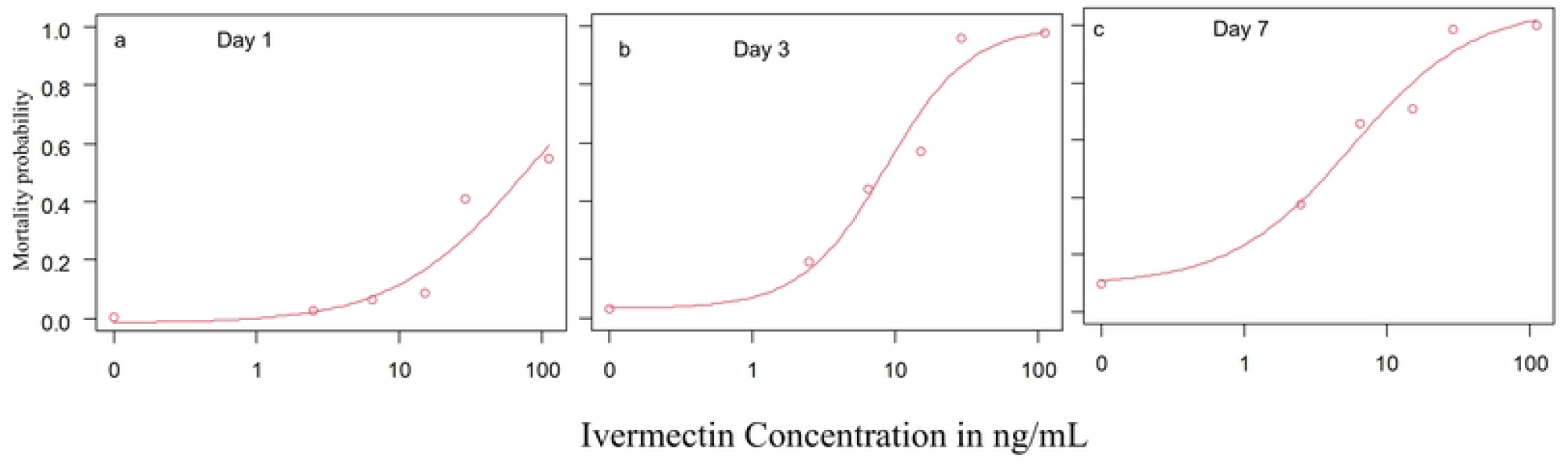
*An. coluzzii* dose-response curves to ivermectin at days 1, 3 and 7.

